# The classic psychedelic DOI induces a persistent desynchronized state in medial prefrontal cortex

**DOI:** 10.1101/2023.02.26.529963

**Authors:** Randall J. Olson, Lowell Bartlett, Alex Sonneborn, Zachary Bretton-Granatoor, Ayesha Firdous, Alexander Z. Harris, Atheir I. Abbas

## Abstract

Administration or consumption of classic psychedelics (CPs) leads to profound changes in experience which are often described as highly novel and meaningful. They have shown substantial promise in treating depressive symptoms and may be therapeutic in other situations. Although research suggests that the therapeutic response is correlated with the intensity of the experience, the neural circuit basis for the alterations in experience caused by CPs requires further study. The medial prefrontal cortex (mPFC), where CPs have been shown to induce rapid, 5-HT_2A_ receptor-dependent structural and neurophysiological changes, is believed to be a key site of action. To investigate the acute neural circuit changes induced by CPs, we recorded single neurons and local field potentials in the mPFC of freely behaving mice after administration of the 5-HT_2A/2C_ receptor-selective CP, 2,5-Dimethoxy-4-iodoamphetamine (DOI). We segregated recordings into active and rest periods in order to examine cortical activity during desynchronized (active) and synchronized (rest) states. We found that DOI induced a robust decrease in low frequency power and decoupled rhythmic activity from neural population dynamics when animals were at rest, attenuating the usual synchronization that occurs during less active behavioral states. DOI also increased broadband gamma power and suppressed activity in fast-spiking neurons in both active and rest periods. Together, these results show that the CP DOI induces persistent desynchronization in mPFC, including during rest when mPFC typically exhibits more synchronized activity. This shift in cortical dynamics may in part underlie the longer-lasting effects of CPs on plasticity, and may be critical to their therapeutic properties.

## Introduction

Classic psychedelics (CPs) induce profound and multifaceted effects on experience after consumption [1]. The resulting experience is often described as “mystical”, and for many is rated as highly meaningful [2,3]. In recent years, there has been rekindled interest in the potential of CPs, particularly psilocybin, as possible therapeutic agents for treatment-resistant depression and other psychiatric disorders. This work has added to longstanding evidence that CPs can chronically alleviate depressive symptomatology in a dose-dependent manner mediated by the intensity of the experience [4–7].

Classic psychedelics exert their main psychoactive effects through interaction with G_q_-coupled 5-HT_2A_ receptors (5-HT_2A_Rs) [8], and pretreatment with the selective 5-HT_2A_R antagonist ketanserin greatly reduces the subjective effects of the CPs psilocybin and LSD [9–11]. Expression of these receptors is enriched in medial prefrontal cortex (mPFC) where roughly half of all excitatory neurons and 20-30% of all GABAergic interneurons express 5-HT_2A_Rs [12–15]. Their activation by both CPs and non-psychedelic 5-HT_2A_R agonists induces a diverse range of downstream molecular actions that do not clearly differentiate CPs from non-psychedelic 5-HT_2A_R agonists [16]. The most consistent long-term effect across compounds seems to be a potent initiation of structural and functional dendritic plasticity [17], which is similarly blocked by application of ketanserin [18,19]. However, the rapid, circuit-level neurophysiological effects which precede this plasticity are less well understood. Thus, investigating how CPs affect neural circuit function might help determine how they acutely modulate experience and aid in identifying key neural activity which confers superior long-lasting therapeutic properties.

Previous *ex vivo* electrophysiology studies in the mPFC have identified complex acute effects of 5-HT_2A_R activation on the intrinsic excitability and synaptic activity of different neuron subtypes. Whole-cell recordings from layer 5 GABAergic neurons have shown that 5-HT_2A_Rs modulate activity most strongly in fast-spiking interneurons [20], the main inhibitory subtype on which they are expressed [21]. More mixed effects were found in pyramidal neurons of rat mPFC, in which 5-HT_2A_Rs were found to either hyperpolarize [22] or depolarize [23–25] membrane potential. Synaptic activity impinging onto mPFC pyramidal neurons is also modified by 5-HT_2A_Rs, which increase the frequency and amplitude of spontaneous and evoked inhibitory [26] and excitatory [27] postsynaptic currents. Furthermore, 5-HT_2A_R activation has been shown to bias these synapses toward a more asynchronous mode of evoked neurotransmitter release [28], which can alter the timing of communication between neurons.

Taken together, these acute *ex vivo* effects of 5-HT_2A_R activation would be expected to alter local field potentials (LFPs), spiking, and synchrony in mPFC circuits *in vivo*. However, to our knowledge only a few studies have investigated these phenomena in freely behaving rodents. In the dorsal anterior cingulate subregion of the mPFC, systemic administration of the 5-HT_2A/2C_-selective CP DOI altered the firing rate of a subset of neurons and decreased gamma power [29]. More ventrally, in the prelimbic and infralimbic cortices, CPs produced the opposite effect on gamma power [30,31], implying subregion-specific mechanisms of CPs in the mPFC. This effect was reported to be significantly modulated by the behavioral state of the animal [30], which suggests a more complex interaction between CPs and cortical activity than previously thought. Studying these behavioral state-dependent effects of CPs on neural network activity is especially intriguing in more ventral mPFC, which is proposed to be analogous to the human subgenual and pregenual ACC [32], two brain regions known to be dysregulated in major depressive disorder [33,34]. Importantly though, neither of the studies in ventral mPFC mentioned above recorded spiking and LFP simultaneously in response to CP administration. Learning how CPs regulate the behavioral state-dependent relationship between spiking and LFP in this region would provide a more complete picture of the complex network effects that may contribute to their therapeutic properties.

To that end, we recorded single units and LFPs in the ventral mPFC of freely behaving mice before and after administration of DOI, with the goal of studying its effect on spiking, brain rhythms, and population activity patterns. We predicted that DOI would lead to a behavioral state-dependent shift in synchronized population dynamics and associated brain rhythms. To assess this, we segregated recordings by behavioral state – active versus rest – and compared pre- and post-DOI mPFC rhythmic power, single neuron firing, and measures of population dynamics. We found that activity in fast-spiking neurons, LFP at multiple frequencies, and the synchrony between neural population dynamics and LFP across a range of frequency bands were all disrupted after administration of DOI in a manner that depended on behavioral state.

## Results

### The CP DOI modulates LFP power in a behavioral state-dependent manner

To examine the interaction between behavioral state and the effect of the CP DOI on cortical activity, we implanted mice with 14 tungsten stereotrodes in the left mPFC. Electrodes were connected to an electronic interface board that facilitated the acquisition of synchronized electrical and video recordings of mPFC neural activity using a Neuralynx acquisition system (see Methods for further details). Mice were placed in a box and allowed to freely explore for a 15-30 minute baseline period, after which they were injected intraperitoneally (i.p.) with 5 mg/kg of DOI or saline. Mice continued free exploration for another 60 min as recordings continued.

We used DeepLabCut [35] to calculate the position and velocity of the animal throughout the recording session to segregate the animal’s behavioral state into active and rest periods (see Methods for details), as LFP is known to be modulated by behavioral state [36–38]. Alignment of velocity traces to wavelet power spectra of mPFC LFP recordings during baseline (Figure 1A) revealed the typical relationship of higher delta (2-4 Hz) and lower theta (6-10 Hz) power during rest states and subsequent reversal of this relationship during active states (Figure 1B). We did not see state-dependent modulation of gamma power (Figure 1C); however, this is not surprising given that our behavioral state classification segregates between lighter rest periods (very low velocity but still interacting with the environment) and active periods. Deeper rest and sleep are associated with decreased gamma power [39–41], but we rarely captured deep rest periods as the mice were mostly engaged during recording. After classifying behavioral state-dependent dynamics, we examined the effect the CP DOI had on these dynamics. DOI-treated animals revealed that the typical increase in low frequency delta (2-4 Hz) power which occurs during rest periods, a hallmark of cortical synchronization [42], is significantly decreased approximately 5 min after DOI injection, but not after saline injection (Figure 2A). Averaging normalized low frequency power spectra across animals showed that delta power was significantly decreased after DOI injection (Figure 2B) but not saline injection. We also found the typical increase in theta power during active states was decreased only after DOI but not saline injection (Supplementary Figure 1A-B). Neither DOI nor saline affected beta power (Supplementary Figure 2C-D).

**Figure 1:**
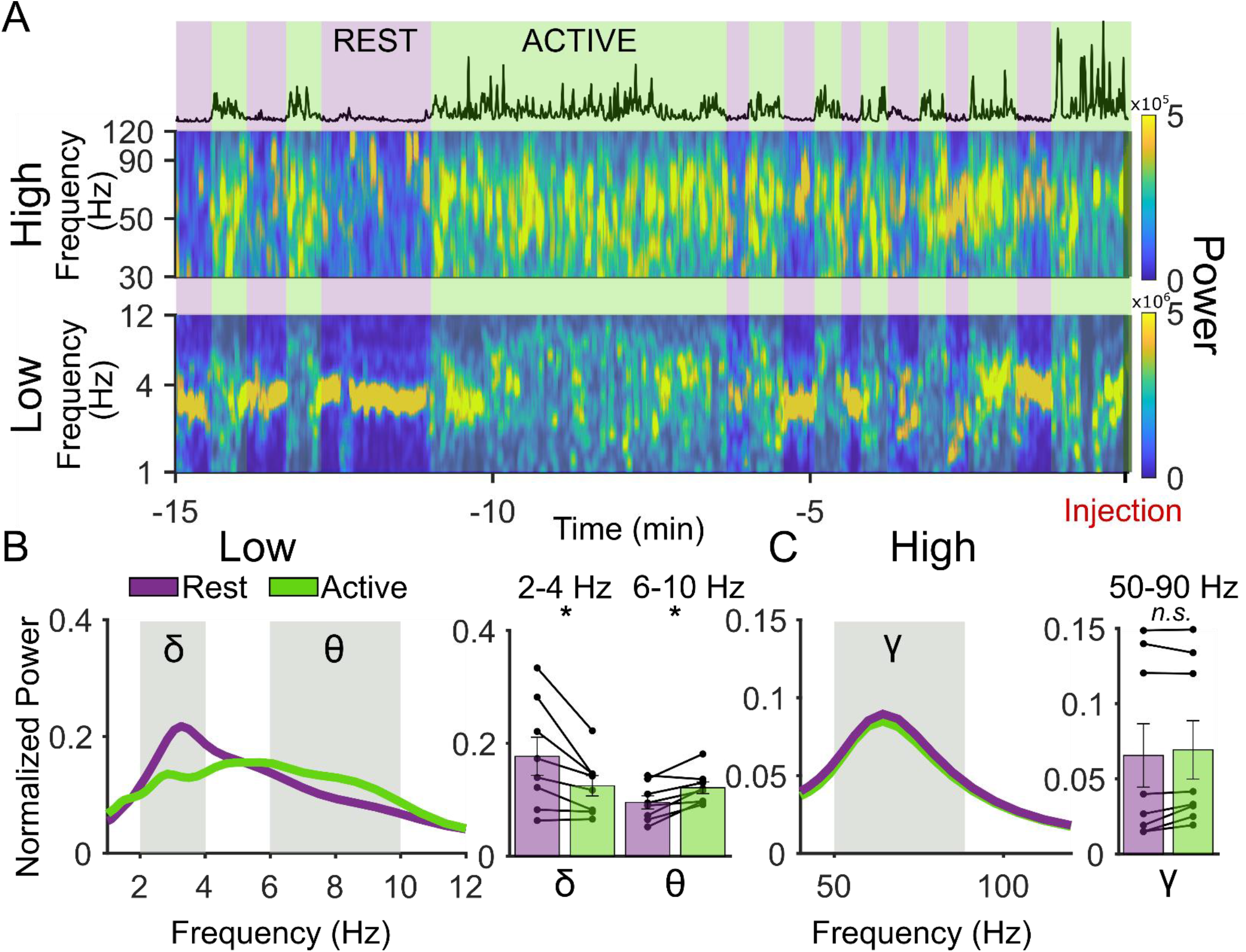
LFP power is behavioral state-dependent. A) Top: Raw power spectrogram from 30-120 Hz of an example animal during baseline (15 min prior to injection (red) of DOI until injection of DOI). Bottom: Same as top but for lower frequencies (1-12 Hz). Black trace above spectrograms is velocity of animal. Purple and green panels denote labeled behavioral state (rest vs. active). B) Left: Average normalized power by frequency for 1-12 Hz for baseline rest and baseline active averaged across all animals shows increased delta power during rest and increased theta during active periods. Right: Average power by animal for baseline rest is greater than baseline active in the delta frequency (2-4 Hz), p=0.0156 and lower for theta frequency (6-10 Hz), p=0.0156. C) Left: Average normalized power by frequency for 40-120 Hz for baseline rest and baseline active averaged across all animals. Right: Average power by animal for baseline rest vs. baseline active in the gamma frequency (50-90 Hz), p=0.1094.

**Figure 2:**
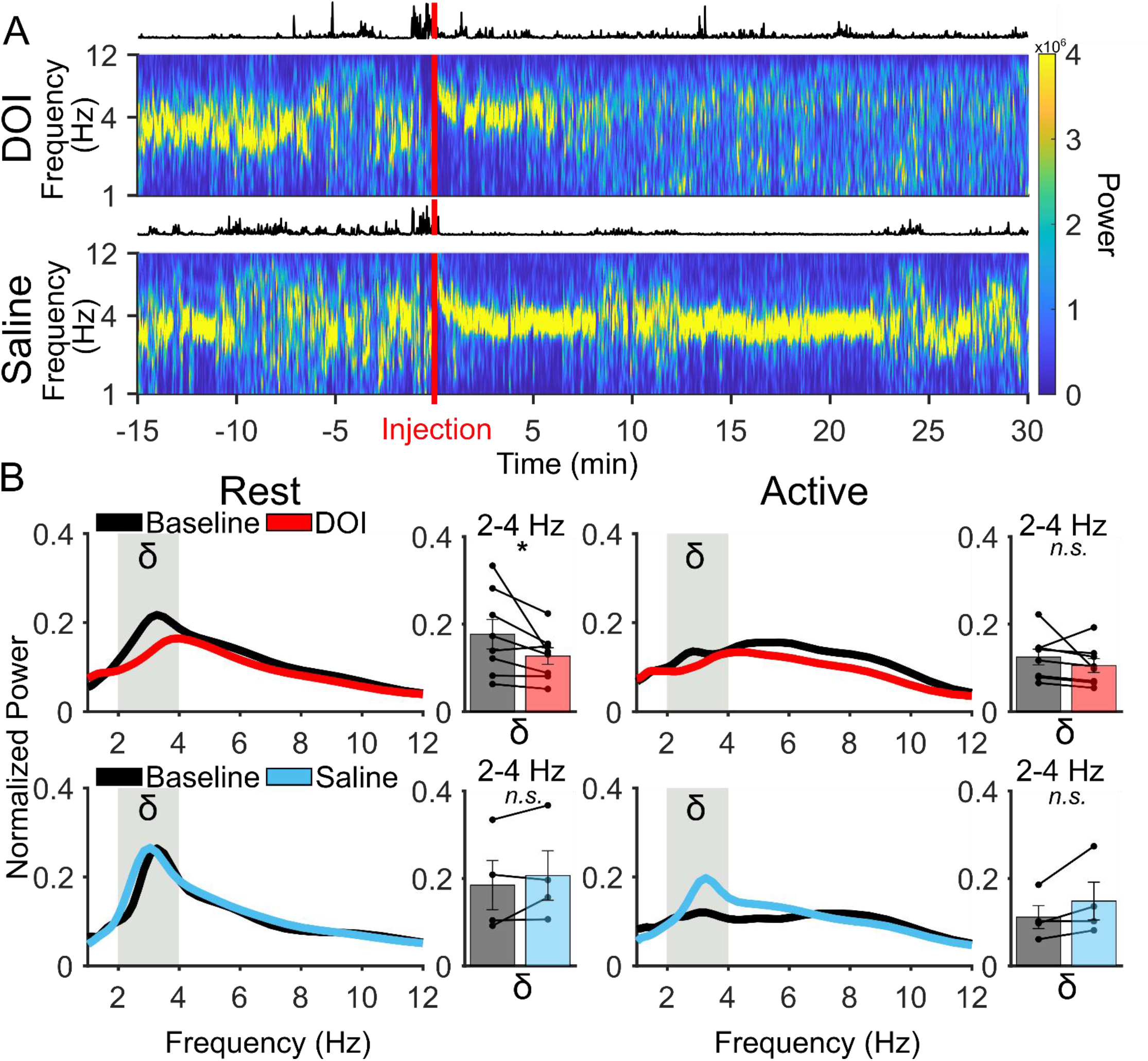
The CP DOI decreases power of low frequency oscillations in mPFC during rest periods. A) Top: Raw power spectrogram from 1-12 Hz of an example animal from 15 min prior to injection (red) of DOI to 30 post Injection of DOI. Bottom: Same as top but with saline. Black traces above spectrograms are velocity of animal. B) Top Left: Average normalized power by frequency (1-12 Hz) for baseline rest and DOI rest averaged across all animals. Average power by animal for baseline rest is lower than DOI rest in the delta frequency (2-4 Hz), p=0.0234. Top Right: Average normalized power by frequency (1-12 Hz) for baseline active and DOI active averaged across all animals. Average power by animal for baseline and DOI active in the delta frequency (2-4 Hz) shows no significant difference, p=0.1484. Bottom Left: Average normalized power by frequency (1-12 Hz) for baseline rest and saline rest averaged across all animals. Average power by animal for baseline and saline rest in the delta frequency (2-4 Hz) shows no significant difference, p=0.3750. Bottom Right: Average normalized power by frequency (1-12 Hz) for baseline active and saline active averaged across all animals. Average power by animal for baseline and saline active in the delta frequency (2-4 Hz) shows no significant difference, p=0.2500.

Alignment of velocity traces with higher frequency spectrograms revealed that systemic injection of DOI led to a significant increase in gamma power (50-90 Hz) in the mPFC (Figure 3A). The DOI-induced increase in gamma power occurred during both rest and active periods (Figure 3B). No increase in gamma power was seen after saline injection (Figure 3B). Taken together, the CP DOI disrupts the typical behavioral state-dependent modulation of oscillatory activity in mPFC.

**Figure 3:**
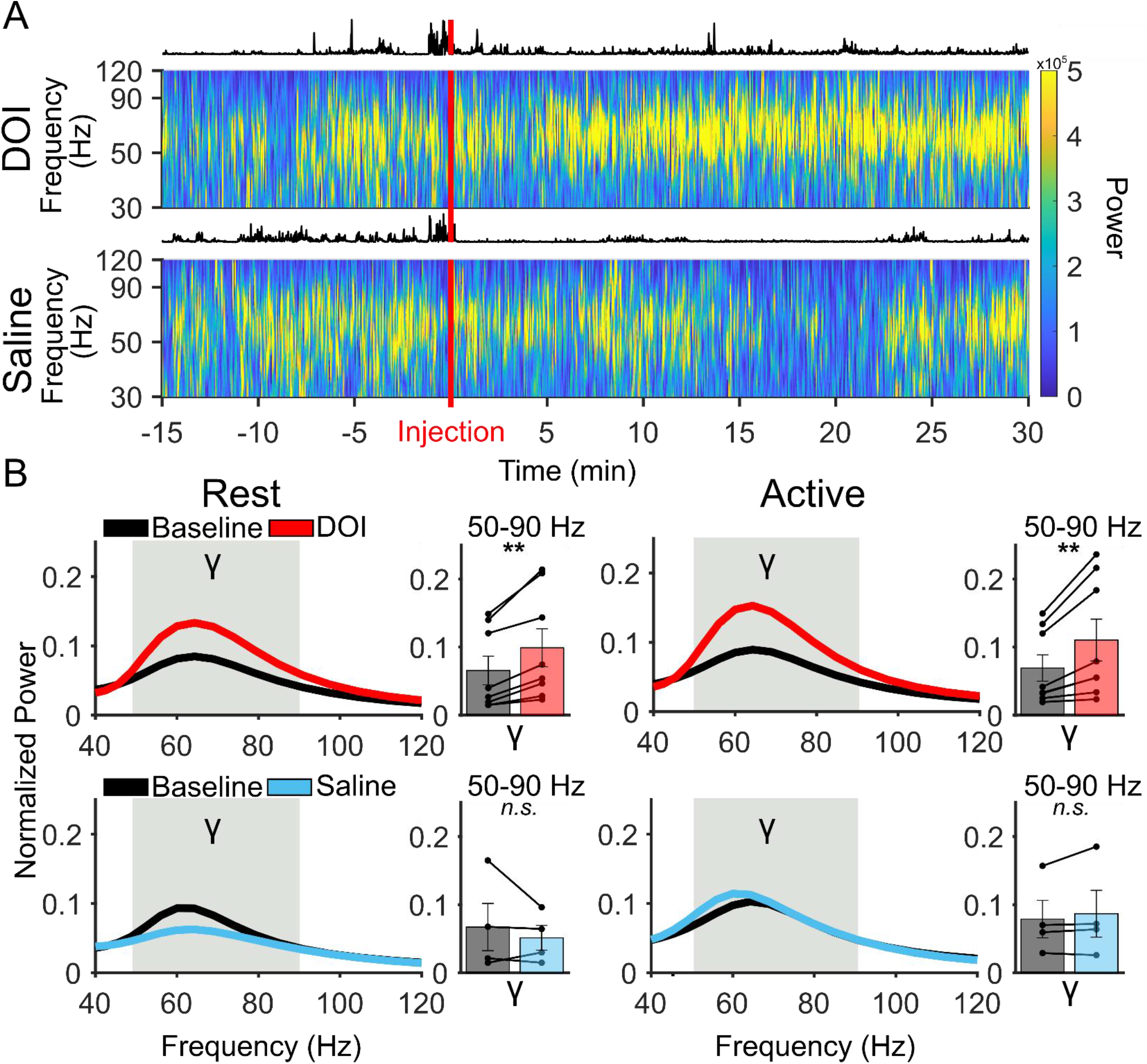
The CP DOI increases power of high frequency oscillations in mPFC irrespective of movement. A) Top: Raw power spectrogram from 30-120 Hz of an example animal from 15 min prior to injection (red) of DOI to 30 post injection of DOI. Bottom: Same as top but with saline. Black traces above spectrograms are velocity of animal. B) Top Left: Average normalized power by frequency (40-120 Hz) for baseline rest and DOI rest averaged across all animals. Average power by animal shows that DOI increases gamma frequency power during rest periods (50-90 Hz), p=0.0078. Top Right: Average normalized power by frequency (40-120 Hz) for baseline active and DOI Active averaged across all animals. Average power by animal shows that DOI increases gamma frequency power during rest periods (50-90 Hz), p=0.0078. Bottom Left: Average normalized power by frequency (40-120 Hz) for baseline rest and saline rest averaged across all animals. Average power by animal for baseline and saline rest in the gamma frequency shows no difference (50-90 Hz), p=0.6250. Bottom Right: Average normalized power by frequency for 50-90 Hz for baseline active and saline active averaged across all animals. Average power by animal for baseline and saline active in the gamma frequency shows no difference (50-90 Hz), p=0.3750.

### DOI preferentially modulates high firing rate neurons

To investigate effects of DOI on the activity of individual neurons in mPFC, we recorded single units before and after injection of DOI or saline and separated neurons into high (>5 Hz; enriched for fast-spiking parvalbumin inhibitory neurons) and low (<5 Hz; enriched for pyramidal neurons) firing rate groups in accordance with their baseline activity. We initially compared average firing rates of these groups during rest and active periods but found that average firing rate did not vary as a function of behavioral state after DOI or saline (Supplemental Figures 2A and 2C). We then sorted them within each group by the magnitude of the change in firing rate after DOI or saline and generated heat-maps of sorted neuron firing rates over the course of the experiment (Figure 4A). The mean firing rate of high firing rate neurons was significantly lower after DOI compared to the baseline period, while there was no significant change in mean firing rate seen with low firing rate neurons (Figure 4B).

**Figure 4:**
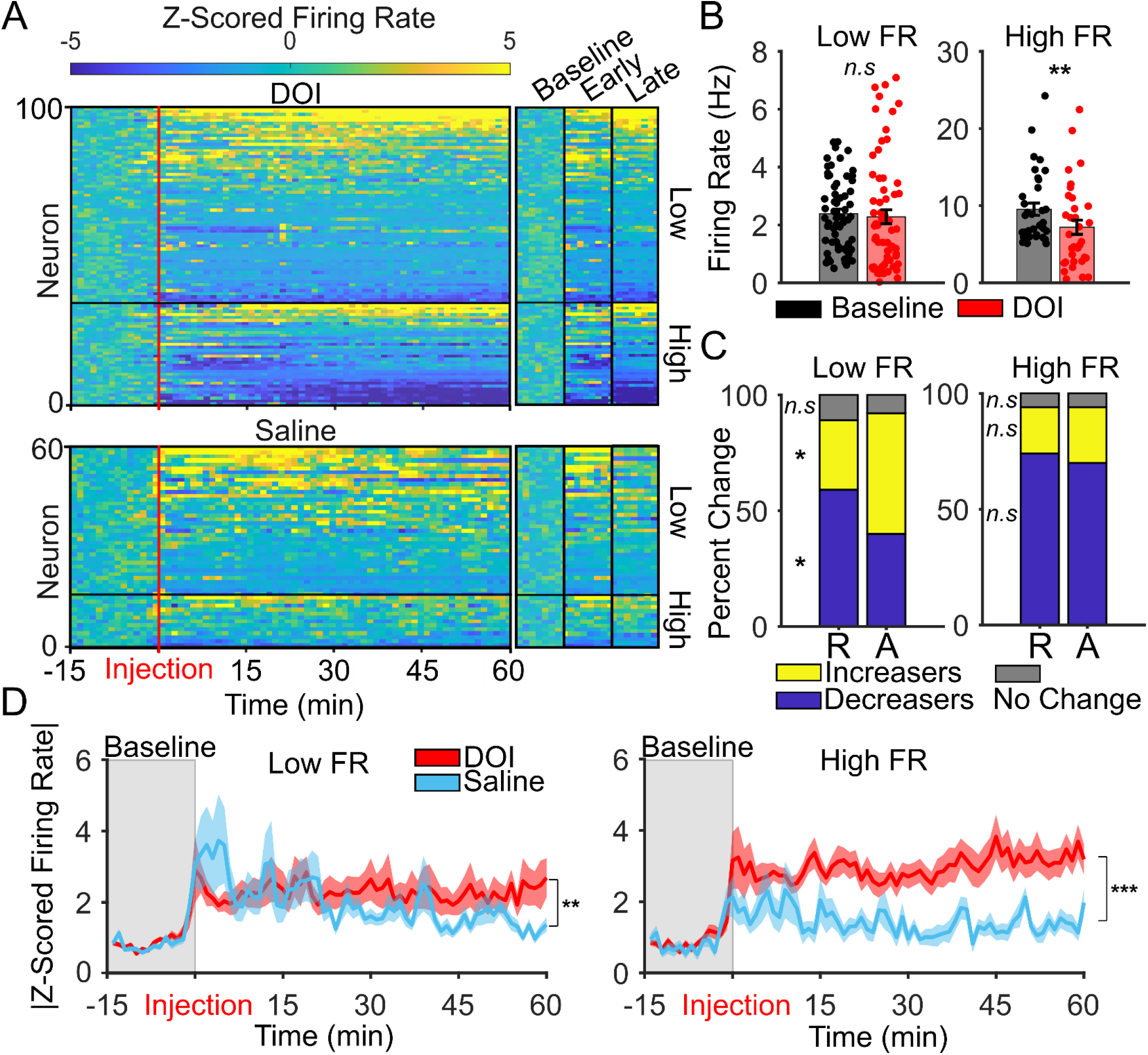
DOI preferentially modulates high firing rate neurons. A) Z-scored firing rate relative to baseline period (−15 min to injection) of all neurons, sorted by firing rate during post injection period, segregated by low firing rate (<5 Hz during baseline) and high firing rate (>5 Hz during baseline) from −15 min prior to injection to 60 min post injection. B) Firing rate changes for high firing and low firing rate neurons between baseline and DOI. Active and rest periods were collapsed as they were not significantly different (Supplemental 2A). Low firing rate neurons showed no difference between baseline and DOI, n.s., p=0.1129. High firing rate neurons showed a significant decrease in DOI from baseline, p=0.0044. C) Percentage of neurons which increase, decrease, or do not change in DOI compared to baseline, separated by firing rate (low vs. high) and by behavioral state (active vs. rest). The percentage of low firing rate increasers are significantly higher in active compared to rest (p=0.0047, by bootstrapping, see methods). The percentage of low firing rate decreasers is significantly higher in rest than active (p=0.0151, by bootstrapping, see methods). The percentage of low firing rate no-change neurons did not change between rest and active, p=0.6259, by bootstrapping, see methods. D) Left: Absolute value of Z-scored firing rate averaged across all low firing rate neurons between DOI and saline, p=0.004. Right: Absolute value of Z-scored firing rate averaged across all high firing rate neurons between DOI and saline, p<0.001.

Breaking the groups down further, we noticed that a larger proportion of high firing rate neurons decreased after DOI as compared to low firing rate neurons (Figure 4C). Interestingly, the relative changes in firing rate for low firing neurons were dependent on behavioral state, showing a significantly higher proportion of cells increasing their firing rate during active periods, suggesting a complex relationship between circuit dynamics and behavioral state (Figure 4C). Accordingly, the magnitude of the change effected by DOI on each neuron, irrespective of direction of change, was significantly larger for high firing rate neurons even after rescaling by Z-scoring (Figure 4D). It is important to note that low firing rate neurons also significantly changed their overall magnitude of firing, however, because the population of low firing neurons is heterogeneous as to direction of firing rate changes, the average firing rate does not change. Overall, DOI has a predominantly inhibitory effect that is larger in terms of the proportion of neurons inhibited and the magnitude of inhibition in high firing rate neurons relative to low firing rate neurons. Furthermore, that effect of DOI on single neuron spiking is modulated by behavioral state, albeit in a complex way.

### DOI disrupts the characteristic relationship between LFP and spiking activity during rest periods

LFPs, which are predominantly a reflection of dendritic input [43], exhibit a characteristic relationship with spiking which depends on a brain region’s afferent and local connectivity [40]. If DOI is indeed disrupting rest-related synchronization within cortex, we predicted that we would see a disruption of that characteristic relationship. To examine this possibility we performed latent dynamical canonical correlation analysis (CCA) [44]. Latent dynamical CCA analysis takes advantage of ongoing neural data and is well-suited for examining the relationship between LFP power and spiking over longer periods of time, unlike phase locking analyses which are better suited for event-centered, trialized data during stereotyped behaviors. During latent dynamical CCA, the dimensionality of population spiking is reduced and a correlation between that reduced signal and bandpass-filtered LFP is calculated to quantify the relationship between the latent dynamics of population spiking and LFP power. Figures 5A-B show an example LFP and the first dimension of the projected population spiking activity before and after DOI for both rest and active periods. In the example, the signals appear less correlated after DOI only during rest. Latent dynamical CCA before and after DOI or saline during both rest and active states revealed that DOI disrupted the correlation between spiking and all frequency bands during rest, and only the theta frequency band during active periods (Figure 5B), as would be expected if DOI induced an active-like desynchronization that persisted into the rest period. In contrast, saline did not affect the canonical correlations between band-specific power and population activity (Supplementary Figure 2E).

**Figure 5:**
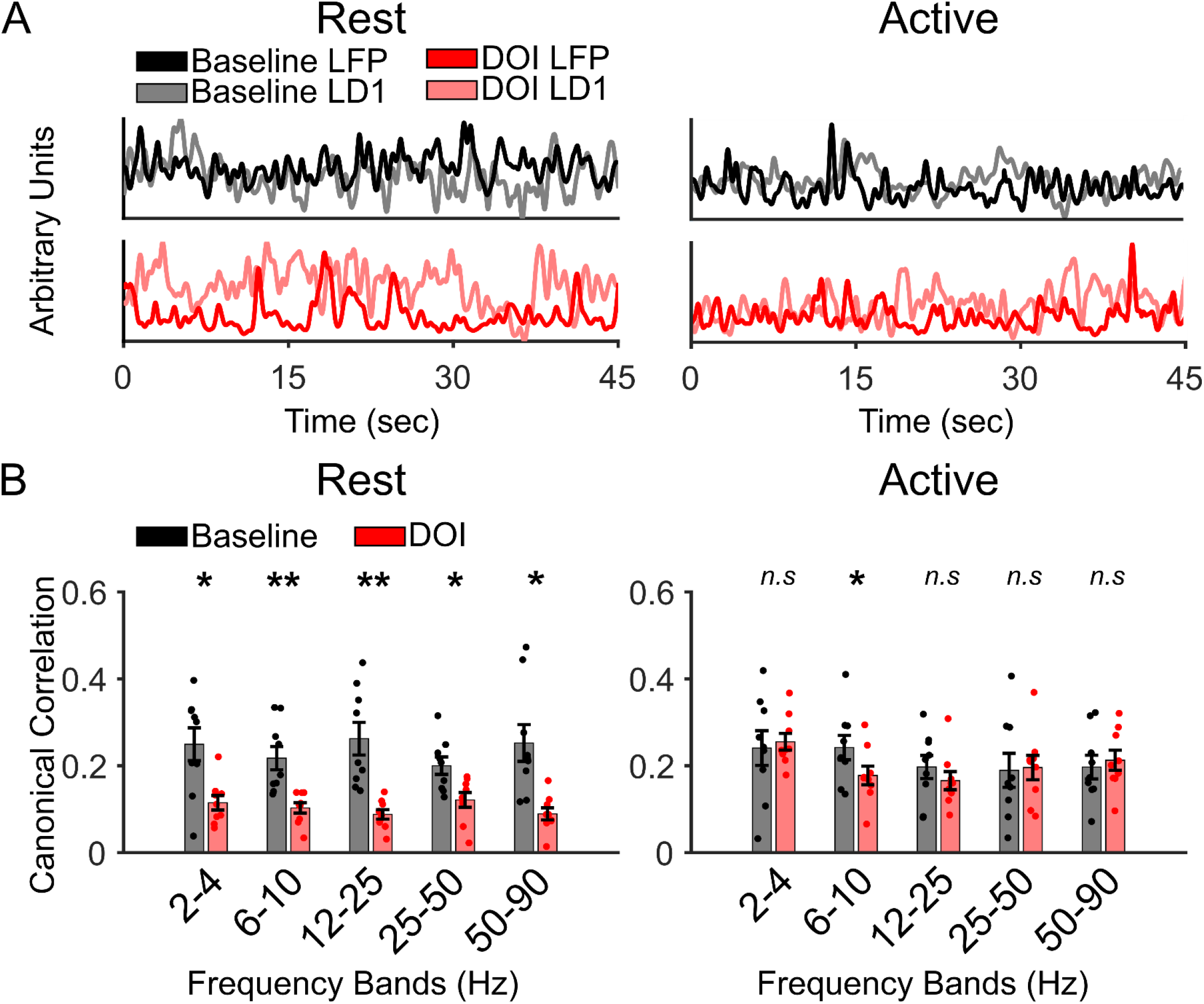
DOI disrupts the characteristic relationship between LFP and spiking activity during rest periods. A) Left: Example traces comparing the first latent population dynamic (LD1) to a single animal’s LFP trace in rest, note the decorrelation seen during DOI. Right: Example traces comparing the first latent population dynamic (LD1) to a single animal’s LFP trace in active. B) Left: Canonical correlation between population latent dynamics of neurons and LFP for each animal by frequency band between baseline and DOI during rest shows the canonical correlation is significantly lower after DOI as compared to baseline across all frequencies (by frequency: p=0.0195, 0.0039, 0.0039, 0.0391, 0.0117). Right: Canonical correlation between population latent dynamics of neurons and LFP for each animal by frequency band between baseline and DOI during active periods shows the canonical correlation is not changed after DOI as compared to baseline across all frequencies except in the theta band (by frequency: p=0.6523, 0.0273, 0.3008, 0.8203, 0.7344).

We also wanted to investigate if spiking patterns changed as a result of decoupling from the LFP during rest that occurs after DOI. To asses this, we calculated the entropy of each neuron’s distribution of inter-spike-intervals. We found that low firing rate neurons exhibited a significant decrease in entropy after DOI compared to baseline, whereas high firing rate neurons were unchanged, albeit trending toward increased entropy (Supplemental Figure 2B). Saline had no effect on spike entropy (Supplementary Figure 2D). Taken together, these results suggest a fundamental change in network dynamics after the CP DOI.

## Discussion

In this study, we recorded mPFC single units and LFP before and after administering the psychedelic DOI during freely moving behavior, which was segregated into rest and active periods. We saw four key circuit-level changes in mPFC after DOI. First, the usual rest-related synchronization that typically occurs in rodents [45], which is associated with an increase in delta power, was attenuated after DOI. Second, we observed an increase in high gamma power (50-90 Hz) irrespective of whether the mouse was active or at rest. Third, there was a significant decrease in the firing rate of fast-spiking, putative parvalbumin inhibitory neurons which also occurs regardless of behavioral state. Fourth, we see a decoupling of mPFC spiking from the LFP during rest. Overall, our findings suggest that DOI induces aberrant desynchronization that persists into rest periods, when brain activity is typically more synchronized.

Our results showing decreased delta power in mPFC during rest agree with other studies using the CP DOI in anesthetized rats [46,47], and with human EEG studies using DMT [48] and psilocybin [49]. DOI also induced a broadband increase in gamma power, another effect reported elsewhere in mice [31]. This is in contrast to Wood et al. [29], who observed a decrease in gamma power. However, their recordings were done in the anterior cingulate cortex, which is known to be functionally and anatomically different from more ventral mPFC [50]. A more recent study in the anterior cingulate cortex, which performed experiments in head fixed mice, similarly found reduced cortical synchrony but a general increase in population firing [51]. Furthermore, our data showing decreased fast-spiking activity after DOI is inconsistent with previously published results showing the opposite [52]. The reason for this discrepancy may be due to the chloral hydrate used to anesthetize animals for recording, which is known to directly affect the serotonergic system [53].

Different behavioral states are canonically characterized by oscillatory activity in distinct frequency bands [45]. In rodent and monkey cortex, rest and quiet wakefulness are normally associated with synchronized, low-frequency oscillations, which transitions to higher-frequency, desynchronized activity during more active engagement with the environment [36,37]. Fast-spiking inhibitory neurons are known to organize the higher-frequency gamma rhythms [54,55], and blocking the firing of these neurons desynchronizes cortical networks and increases broadband gamma power [56]. Therefore, it is plausible that the aberrant desynchronized activity that we see during rest periods is partly due to DOI disrupting the dynamics of fast-spiking neurons, which prevents the normal transition to more synchronized brain states during rest/quiet wakefulness.

More active cortical states with decreased delta power and increased gamma power normally reflect a shift toward more local computations [39,57]. These enhanced gamma power states are theorized to be optimal for inducing synapse-specific long-term plasticity [58], the precision of which relies on feed-forward inhibition involving parvalbumin positive, fast-spiking interneurons [59]. Since DOI seems to be modulating these neurons, the observed desynchronized rest state could lead to long-lasting changes in associative plasticity [60], which may underlie some of the experiential or therapeutic effects of CPs. This desynchronized state induced by DOI is consistent with the theory that psychedelics effectively enable neural networks to escape their strongest previously entrained patterns of activity, leading to a more labile and dynamic state [61,62].

As a precedent for this idea, several studies have shown long-lasting 5-HT_2A_R-dependent plasticity following CP administration. An *in vitro* study showed that when exposed to CPs, rat cortical neurons displayed an increase in dendritic arbor complexity, dendritic spine growth, and new functional synapse formation [18]. *In vivo*, a single dose of psilocybin resulted in dendritic growth and synaptic activity in layer 5 apical dendrites of the mouse mPFC [19]. Both of these findings persisted for at least 24 hours after drug removal and were dependent on 5-HT_2A_Rs. Slice studies report a more rapid effect of 5-HT_2A_R activation in which DOI strengthens NMDA currents but weakens AMPA currents within minutes, while gating the induction of spike-timing dependent depression in mPFC at thalamocortical synapses [63,64]. These long-lasting neuroplastic changes induced by CPs may be caused at least in part by the rapid circuit changes occurring after CP administration reported in this article.

It is apparent that the circuit level effects of DOI are complex, and our results highlight the need for future *in vivo* work on this subject in awake, freely behaving animals [65]. Studying the effect of DOI during behaviors that depend on circuits and cell subpopulations expressing 5-HT_2A_ receptors will be particularly valuable. The effect of DOI on high broadband gamma is especially intriguing for future study, as the same phenomenon is seen in humans with ketamine [66] and other CPs like psilocybin [67]. This could be related to the known effectiveness of ketamine and several CPs for alleviating treatment-resistant depression, which supports the idea that elevated broadband gamma power in the mPFC may be an early translational neurophysiological correlate of antidepressant efficacy. Follow up work will require more detailed experiments to definitively determine which neuronal subpopulations are most affected by DOI and explore whether these effects are being driven predominantly locally or due to effects on one or more long range inputs.

Overall, our results suggest that classic psychedelics like DOI induce rapid circuit level changes in the mPFC. These circuit changes fundamentally alter the typical dynamics associated with different behavioral states, favoring persistent cortical desynchronization. This shift toward desynchronization may be a correlate of the experiential effects of psychedelics and of a more labile neural state which drives subsequent neuroplastic changes observed after psychedelic administration.

## Methods

### Animals

All procedures were conducted in accordance with the US NIH Guide for the Care and Use of Laboratory Animals and approved by the New York State Psychiatric Institute Institutional Animal Care and Use Committee at Columbia University. Eight adult (age 12-24 weeks) C57BL/6J (Jackson Labs, stock number 000664) experimental mice were used for DOI experiments, while a subset of these mice (n = 4) received saline injections one week following DOI administration and were used for all saline analyses.

### Surgical procedures

Mice were anesthetized with 1%–3% vaporized isoflurane in oxygen (1 L/min) and placed in a stereotaxic apparatus. A craniotomy was made to allow for implantation of a 28 microwire bundle (14 stereotrodes; 13 micron tungsten wire, California Fine Wire) implanted in left mPFC (−0.35 ML, +1.85 AP, 1.3 below brain surface). A ground screw was placed over the cerebellum and a reference screw was placed over the orbitofrontal cortex. Electrodes were connected to a 32-channel Omnetics electrode interface board using gold pins (Neuralynx). Electrode placements were confirmed using an electrolytic lesion (5 mA, 10 s). Mice were allowed to recover for one week post-surgery before behavioral testing and were monitored closely during recovery.

### Behavior

Animals were recorded in a novel open field environment and were allowed to move freely. After 15 minutes of baseline recording mice received a 5 mg/kg i.p. injection of racemic 2,5-dimethoxy-4-iodoamphetamine (DOI) and then placed back in the open field environment to freely behave for 60 min. A subset of animals (n=4) were exposed to the same experiment as described above except that saline was injected.

### Neural recording

A Digital Lynx system (Neuralynx, Bozeman, MT) was used to amplify, band-pass filter (1-1000 Hz for LFPs and 600-6000 Hz for spikes), and digitize the electrode recordings. LFP sampling rates were 2 kHz and spikes were collected at 32 kHz. Single units were clustered based on the first two principal components (peak and energy) from each channel using Klustakwik (Ken Harris) and visualized in SpikeSorter3D (Neuralynx). Clusters were then visually inspected and included or eliminated based on waveform appearance, inter-spike interval distribution, isolation distance, and L-ratio.

### Segregation by behavioral state

All data analyses was done using custom scripts in MATLAB (Mathworks) unless otherwise identified. Video was recorded at 30 fps and synchronized to neural data (Digital Lynx). Videos were then imported into DeepLabCut (DLC, https://github.com/DeepLabCut), and positional markers were manually set for head, body and tail for 200 randomly selected frames throughout the entire recording. DLC was then run and average position data from the head, body and tail was calculated per frame for each animal. Due to the imperfect nature of DLC and errors in position, data was cleaned by removing any points that were outside of the physical space the animal could move, and those values were filled by the previous value. Velocity outliers were further removed via a hampel identifier and smoothed via a moving window of 5 fps. Each velocity trace was then synchronized to the existing video and manually inspected. Upon further inspection, large outliers that were not identified by the hampel identifier were removed using a threshold value of 120-fold greater than the threshold value for active/rest. The active/rest threshold velocity value was identified base on visual inspection of velocity trace and video agreement across all animals, and one threshold for all animals was assigned. This threshold was then applied to the velocity trace resulting in a binary vector, 0 denoting rest and 1 denoting active. This vector still resulted in 0 or 1 periods that only last a few frames. These micro rest/active states were too small to perform further analyses on. As a result, sequential 5 second windows were used to identify the consensus of 0s or 1s in each window, changing to either all zeros or all ones for a given window.

### Local field potential analyses

#### Power

LFP data was segregated for the baseline window (−15 min to −5 min prior to DOI or saline injection) and for the experimental window (DOI or saline, +25 min to +60 min after injection). For both the baseline window and the experimental window, bins corresponding to the “active” and “rest” periods were segregated and respectively concatenated into 4 conditions, (1) baseline active, (2) baseline rest, (3) experimental active, and (4) experimental rest, where experimental refers to DOI or saline. Each condition was normalized by the root mean square of the whole signal, and the analytic signal was calculated via the continuous 1-D wavelet transform (CWT using Morse wavelet) with frequency limits 1-120 Hz, symmetry parameter gamma (*γ*) = 3, time-bandwidth product equal to 60, and 10 voices per octave. CWT uses L1 normalization. We calculated the power of the LFP by taking the square of the absolute value of the analytic signal. The power was then segregated into canonical frequency bands: delta (2-4 Hz), theta (6-10 Hz), beta (12-25 Hz), low gamma (30-50 Hz) and high gamma (50-90 Hz). For spectrograms in Figures 1, 2 and 3 the entire non-segregated signal was used with which power was calculated as described above, with the smoothed non-binarized velocity trace. For quantification of LFP each animal’s power was averaged across conditions and then compared using Wilcoxon signed-rank tests.

### Single unit analyses

#### Firing Rate

Spike times sampled at 32 kHz were imported into MATLAB, each neuron having an associated spike train with spike times given to the nearest nanosecond. Spikes were binned in 1 ms bins, and we calculated the baseline firing rate in Hz using the same LFP baseline window. Neurons with baseline firing rates below 0.5 Hz were omitted from further analysis. The remaining neurons were classified as either “high” or “low” firing rate neurons based on their average baseline firing rates, with 5 Hz being the cutoff between the two groups. Average firing rate for each of the same four conditions used in the LFP (1) baseline active, (2) baseline rest, (3) experimental active, and (4) experimental rest was calculated, where experimental refers to DOI or saline. Wilcoxon signed-rank tests were used to compare firing rates of neurons across conditions.

#### Firing Rate Changes

A wide range of responses to drug administration were observed in neuronal activity in the form of firing rate changes. To categorize neurons as increasing their firing rate relative to baseline or decreasing their firing rate relative to baseline, referred to as “increasers” or “decreasers,” spike trains were bootstrapped to ensure the observed change in activity was statistically significant. Bins from the baseline and experimental period were concatenated into a single spike train, which was used to generate 1,000 shuffled spike trains by randomly selecting a number of bins equivalent to either the length of the baseline spike train or the length of the experimental spike train to create a shuffled baseline or experimental spike train, respectively. The difference in these spike trains was calculated for each pass, and the observed difference was compared to the difference of the collection of shuffled spike trains. If the observed difference was greater or less than 97.5% of the shuffled differences, the cell was labeled as an increaser or decreaser, respectively. Neurons not significantly different from the shuffled spike train were labeled as no change. To test proportions of increasers and decreasers across conditions we bootstrapped the proportions 10,000 times.

#### Absolute Z-Scored Firing Rate

To assess the firing rate dynamics in experimental conditions and examine firing rate changes from baseline, spikes were binned into 60 second bins. Firing rates in each bin for each neuron were Z-scored relative to the neuron’s mean firing rate across the 10 minute baseline period. Z-scored firing rates were averaged separately across the population of low- and high-firing rate neurons, respectively, and then the absolute value was taken. Average absolute value Z-scored firing rates after DOI were then compared against firing rates after saline for both populations of neurons using Wilcoxon signed-rank tests.

#### Spike Entropy

Spike times sampled at 32 kHz were imported into MATLAB, each neuron having an associated spike train with spike times given to the nearest nanosecond, and then converted to spike times in seconds. The inter-spike-interval (ISI) distribution for each neuron was calculated using MATLAB’s built in histogram function with 50 bins at a width of 5 milliseconds. A probability mass function (PMF) was then calculated by dividing the count in each bin by the sum of all counts, and probabilities of zero were excluded. The normalized Shannon entropy, *h*, was calculated from the PMF. Wilcoxon signed-rank tests were used to compare entropy of neurons across conditions.

### Local field potential and population analysis

#### Latent Dynamics

To analyze how local field potentials influence ongoing mPFC spiking population dynamics, we examined the correlation between latent population activity and LFP. For a more in-depth review of this analysis see Gallego-Carracedo et al. (2022) [44]. First LFP power in the mPFC was calculated as stated in our LFP methods above. Spikes and LFP power were binned in 30 millisecond bins and smoothed with a Gaussian kernel with a width of 50 milliseconds. The firing rate was square-root transformed. This yields an n by T matrix, where n = number of neurons, and T = time points. A low dimensional manifold of this matrix was made by performing PCA and taking the first 10 PCs, yielding a matrix X. The smoothed firing rates of n neurons were then projected onto the low dimensional manifold matrix X created with the PCA, yielding an m by T latent manifold L, where m is the dimension of the manifold. Finally, canonical correlation analysis was performed between the latent manifold L and the LFP power at a given frequency band, yielding correlation values from 0 to 1. This analysis was performed for each condition (1) baseline active, (2) baseline rest, (3) experimental active, and (4) experimental rest, where experimental refers to DOI or saline. Wilcoxon signed-rank tests were used to compare the canonical correlations that resulted from the latent dynamical CCA across conditions for each frequency band.

