## Supplemental Figures for "The classic psychedelic DOI induces a persistent desynchronized state in medial prefrontal cortex"

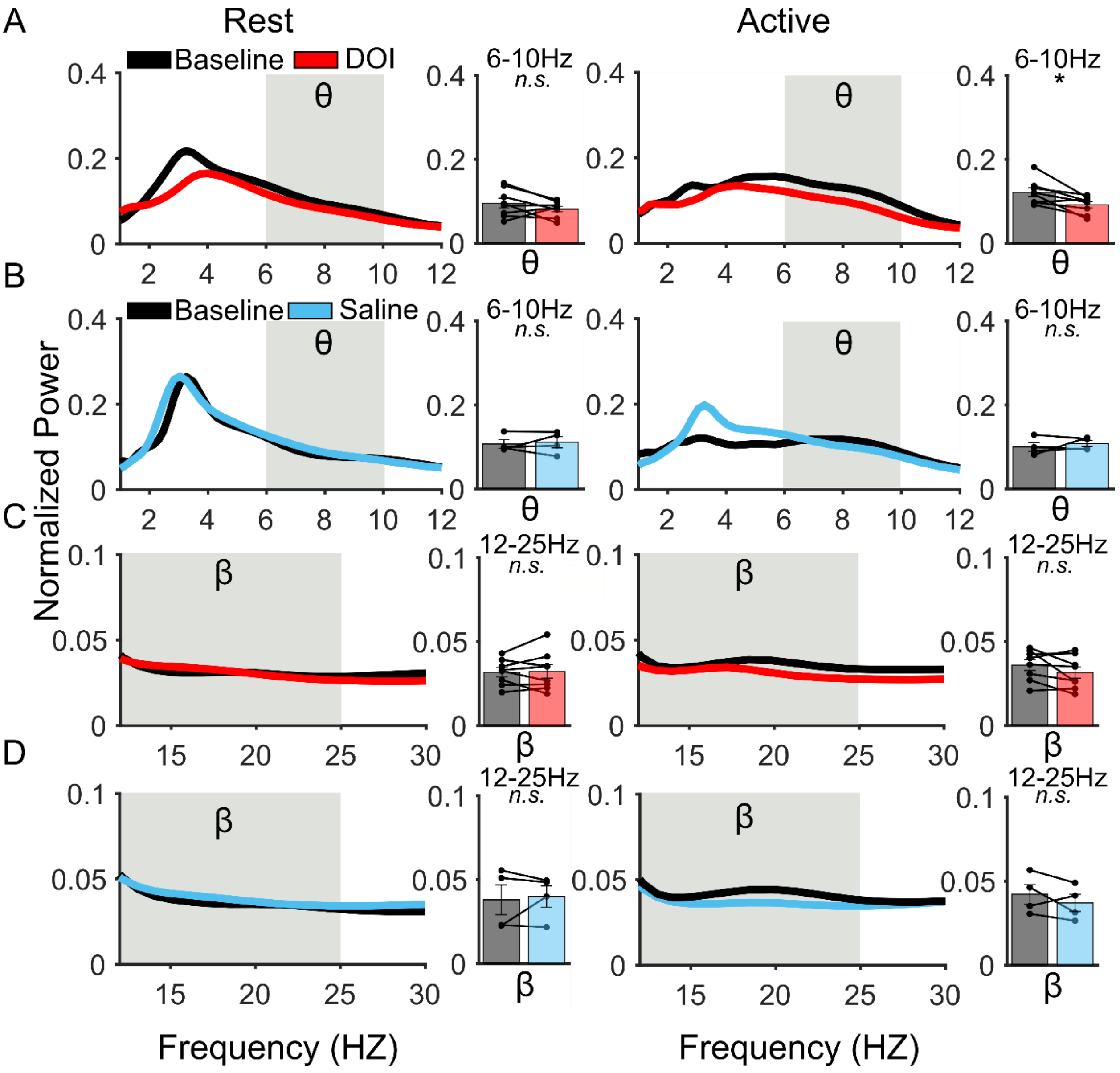


<u>Supplemental Figure 1: DOI decreases LFP theta power only during active periods and beta LFP power remains unchanged.</u> A) Left: Average normalized power by frequency (1-12 Hz) for baseline rest and DOI rest averaged across all animals, then quantified for the theta frequency band (6-10 Hz) n.s, p=0.3125. Right: Average normalized power by frequency (1-12 Hz) for baseline active and DOI active averaged across all animals, then quantified for the theta frequency band (6-10 Hz) shows that DOI decreases theta during active periods, p=0.0156. B) Left: Average normalized power by frequency (1-12 Hz) for baseline rest and saline rest averaged across all animals, then quantified for the theta frequency band (6-10 Hz) n.s., p=0.8750. Right: Average normalized power by frequency (1-12 Hz ) for baseline active and saline active averaged across all animals, then quantified for the theta frequency band (6-10 Hz) n.s., p=0.8750. C) Left: Average normalized power by frequency (12-30 Hz) for baseline rest and DOI rest averaged across all animals, then quantified for the beta frequency band (12-25 Hz) n.s., p=0.8438. Right: Average normalized power by frequency (12-30 Hz) for baseline active and DOI active averaged across all animals, then quantified for the beta frequency band (12-25 Hz), n.s., p=0.2500. D) Left: Average normalized power by frequency (12-30 Hz) for

baseline rest and saline rest averaged across all animals, then quantified for the beta frequency (12-25 Hz) n.s., p=0.8750. Right: Average normalized power by frequency (12-30 Hz) for baseline active and saline active averaged across all animals, then quantified for the beta frequency (12-25 Hz), n.s., p=0.3750.

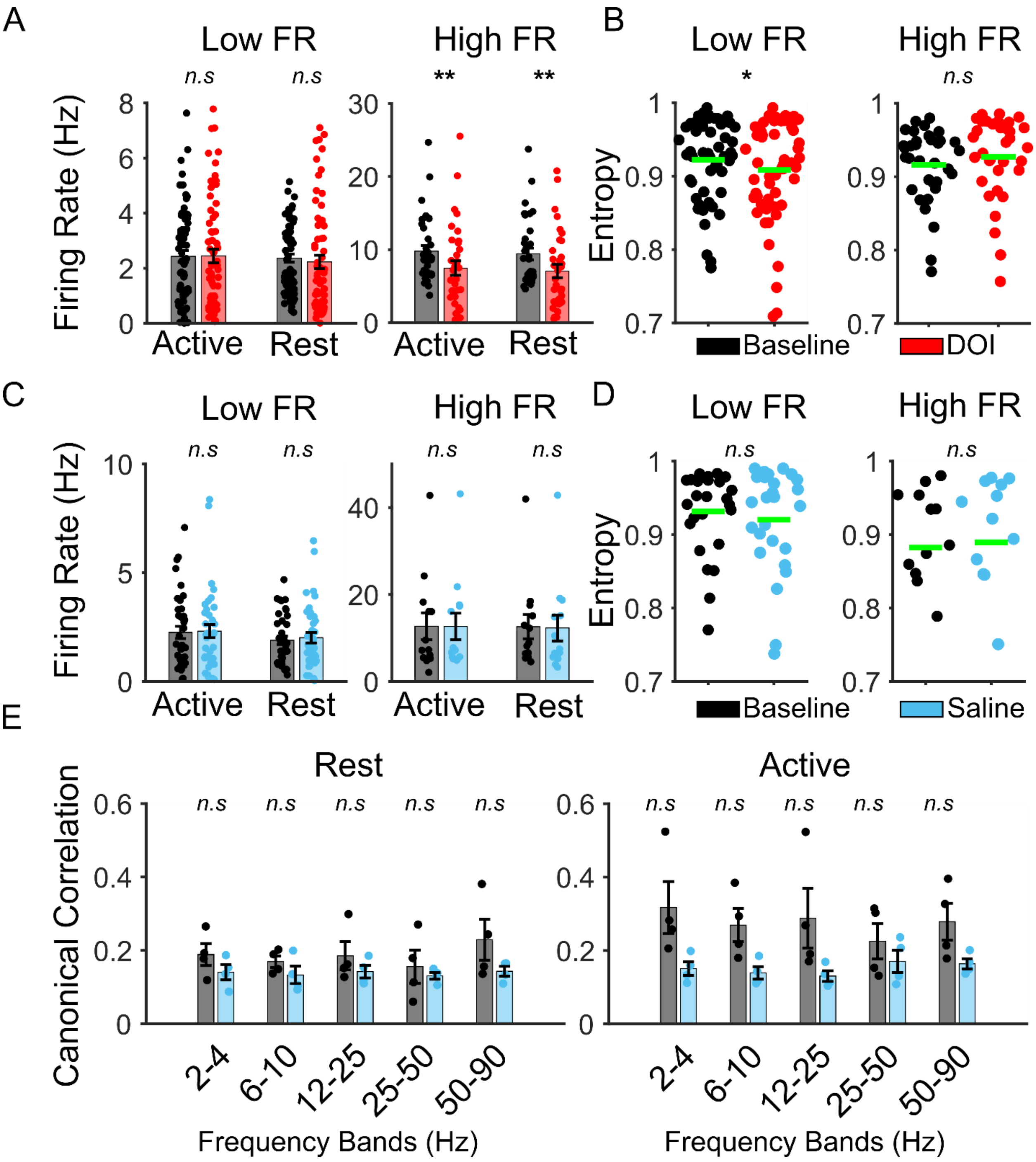


<u>Supplemental Figure 2: DOI preferentially modulates high firing rate neurons and imparts an inverse entropic relationship between low and high firing rate neurons.</u> A) Firing rate changes for high firing and low firing rate neurons between baseline and DOI, segregated by behavior state. High firing rate neurons were significantly decreased in DOI as compared to baseline in both active (p=0.0052) and rest (p=0.0042). Low firing rate neurons in DOI did not change on average for either active (p=0.9975) or rest (p=0.1067). B) Entropy of high firing rate neurons did not change in DOI compared to baseline, p=0.0889. Entropy of low firing rate neurons significantly decreased in DOI compared to baseline, p=0.0158 C) Firing rate changes for high firing and low firing rate neurons between baseline and saline segregated by behavior state.

Neither high firing rate neurons nor low firing rate neurons significantly changed during saline – high active ($p=0.9460$) and high rest ($p=0.9760$), low active ($p=0.8334$) and low rest ($p=0.6478$). D) Entropy of high firing rate neurons did not change in saline compared to baseline, $p=0.8093$. Entropy of low firing rate neurons did not change in saline compared to baseline, $p=0.1909$. E) Left: Canonical correlation between latent dynamics of population spiking and LFP for each animal by frequency band between baseline and saline during rest periods. The canonical correlation is not changed in saline as compared to baseline across all frequencies (by frequency: $p=0.1250$, 0.1250, 0.1253, 0.3750, 0.2500). Right: Canonical correlation between latent dynamics of population spiking and LFP for each animal by frequency band between baseline and saline during active periods, the canonical correlation is not changed in saline as compared to baseline across all frequencies (by frequency: $p=0.1250$, 0.1250, 0.1250, 0.3750, 0.2500).
